# Selective Vulnerability of Tripartite Synapses in Amyotrophic Lateral Sclerosis

**DOI:** 10.1101/2021.10.28.465435

**Authors:** MJ Broadhead, C Bonthron, J Waddington, WV Smith, MF Lopez, S Burley, J Valli, F Zhu, NH Komiyama, C Smith, SGN Grant, GB Miles

## Abstract

Amyotrophic Lateral Sclerosis (ALS) is a fatal neurodegenerative disorder. Separate lines of evidence suggest that synapses and astrocytes play a role in the pathological mechanisms underlying ALS. Given that astrocytes make specialised contacts with some synapses, called tripartite synapses, we hypothesise that tripartite synapses could act as the fulcrum of disease in ALS. To test this hypothesis, we have performed an extensive microscopy-based investigation of synapses and tripartite synapses in the spinal cord of ALS model mice and post-mortem human tissue from ALS cases. We reveal widescale synaptic changes at the early symptomatic stages of the SOD1^G93a^ mouse model. Super-resolution microscopy reveals that large complex postsynaptic structures are lost in ALS mice. Most surprisingly, tripartite synapses are selectively lost while non-tripartite synapses remain in equal number to healthy controls. Finally, we also observe a similar selective loss of tripartite synapses in human post-mortem ALS spinal cords. From these data we conclude that tripartite synaptopathy is a key hallmark of ALS.

## Introduction

Amyotrophic Lateral Sclerosis (ALS) is a devastating form of motor neuron disease, characterised by loss of motor neurons (MNs) in the brain, brainstem and spinal cord, leading to progressive decline of motor control and fatal paralysis, typically within a few years of diagnosis (Zarei et al., 2015). The pathological mechanisms that precede MN cell death remain unclear. One line of evidence suggests that ALS, like many neurodegenerative disorders, could be caused by a loss or a change in the synapses between neurons within the central nervous system (CNS) prior to cell loss (Fogarty, 2019; Gillingwater & Wishart, 2013; Van Zundert et al., 2008; Wishart et al., 2006). Pre-symptomatic and early-symptomatic changes in cortical dendritic spines, neuromuscular junctions and the synaptic inputs to spinal cord MNs have all been reported (Fogarty et al., 2015, 2016; Perkins et al., 2021; Sasaki & Iwata, 1995; Starr & Sattler, 2018; Tremblay et al., 2017; Van Zundert et al., 2008). Separate evidence indicates that astrocytes, the supportive glial cells of the CNS, may be responsible for non-cell autonomous degeneration of MNs (Devlin et al., 2015; Forsberg et al., 2011; Valori et al., 2014; Yamanaka et al., 2008; Yamanaka & Komine, 2018; Zhao et al., 2020). Astrocytes make specialised connections with chemical synapses, forming tripartite synapses, through which they help regulate synaptic structure and transmission (Araque et al., 1999; Perea et al., 2009). Despite evidence that glial cells may contribute to synaptic degeneration in neurodegenerative diseases (Boillée et al., 2006; Henstridge et al., 2019; Tong et al., 2013), it is not known whether tripartite synapses are affected in ALS. We hypothesise that both synaptic and astrocytic dysfunction could be mechanistically linked, and that the tripartite synapse could act as a vulnerable fulcrum of disease in ALS.

To investigate how synapses and tripartite synapses are affected in ALS, we have performed a thorough, quantitative microscopy-based study of millions of excitatory synapses and markers of perisynaptic astrocytic processes (PAPs) in the spinal cord. This was conducted in mice with ALS-causing mutations and in post-mortem spinal cord tissue from human ALS patients. Using high-resolution and super-resolution microscopy, in combination with genetic and immunohistochemical labelling strategies for visualising synapses and astrocytes, our findings reveal that tripartite synapses are selectively vulnerable to degeneration in ALS. This finding suggests that synaptic loss may be mechanistically linked to astrocytic pathology in ALS, and the identification of this vulnerable synapse subtype could pave the way for targeted therapeutic strategies.

## Results

### Widescale Mapping of Mouse Spinal Cord Reveals Synaptic Changes in ALS

To visualise and quantify synaptic changes in ALS mice using a standardised and robust approach, we cross-bred PSD95-eGFP^+/+^ mice (Broadhead et al., 2016; Zhu et al., 2018) with SOD1^G93a^ mice (Gurney et al., 1994), WTSOD1 mice (Gurney et al., 1994) and C9orf72 mice (Liu et al., 2016). The use of PSD95-eGFP mice ensures that all mice expressed fluorescently tagged postsynaptic density (PSD) scaffolding molecule, PSD95, at virtually all excitatory synapses throughout the nervous system (Zhu et al., 2018). As sex-dependent differences in phenotypes have been observed in both the SOD1^G93a^ and C9orf72 mice (Cacabelos et al., 2016; Heiman-Patterson et al., 2005; Liu et al., 2016), males and females were analysed separately.

Consistent with previous reports (Mead et al., 2011), our SOD1^G93a^ mice and PSD95-eGFP x SOD1^G93a^ mice presented with hind-limb tremor and reduced hind-limb splay by approximately 75 days of age (73.6 days in males, 78.2 days in females, t(16)=2.0, p=0.061); see methods). At the point of termination, none of the 8 week SOD1^G93a^ mice displayed hind limb tremor (5 males, 5 females); all the 12 week male (5/5) and some of the 12 week female (3/5) SOD1^G93a^ mice displayed hind limb tremor, and all the 16 week mice SOD1^G93a^ (6 males, 5 females) displayed hind limb tremor. 16 week old SOD1^G93a^ mice were also lower in weight compared to non-Tg controls (males: t(9)=5.2, p=0.0006; females: (t(8)=5.3, p=0.0007). No signs of motor dysfunction or weight loss were observed in WTSOD1 mice up to 16 weeks. Similarly, no signs of acute motor dysfunction or weight loss were seen in C9orf72 mice up to 22 weeks C9orf72 mice did, however, display G4C2 RNA foci, a key molecular hallmark of disease, in the nuclei of spinal cord cells (SI. Fig. 1A-B) and primary cultured astrocytes (SI. Fig. 1C-D).

Tiled, high-resolution maps of PSD95-eGFP synapses were obtained across hemisects of spinal cords which we delineated into spinal laminae I-II, III-IV, V, VI, VII, VIII, IX and X. Each map of a mouse hemi-spinal cord yielded data on approximately 40,000 to 50,000 PSDs. In total, spinal cord synapse maps were obtained from 122 mice, from which over 5 million PSDs were quantified. In this study, our analysis focuses on PSD density (the number per unit area) and PSD size (area). Overall, inter-regional mapping of PSD95 revealed anatomical diversity in synapse number and size comparable to our previous observations (Broadhead et al., 2020), with little difference associated with age or sex.

In both male and female SOD1^G93a^ and non-Tg control mice, synaptic maps were generated at 8 weeks (pre-symptomatic), 12 weeks (early-symptomatic) and 16 weeks of age (early-symptomatic with significant weight loss) (Fig. 1A-B). At 8 and 12 weeks of age there was no difference between SOD1^G93a^ and non-Tg controls in PSD density (Fig. 1C) or PSD size (Fig. 1D). At 16 weeks of age, however, SOD1^G93a^ males showed significantly reduced PSD density (F(1,72)=9.603, P=0.0028) and PSD size (F(1,72)= 15.77, P=0.0002) compared to non-Tg controls. These effects were also significant when SOD1^G93a^ males were compared with both the non-Tg controls and WTSOD1 controls (PSD density: F (2, 104) = 7.493, P=0.0009; PSD size: F (2, 104) = 11.02, P<0.0001). These synaptic changes were widespread across the spinal cord, with the exception of the dorsal laminae I-II; while post-hoc Tukey’s testing revealed that PSD density and size were most significantly affected in lamina VI and VII respectively.

**Figure 1.**
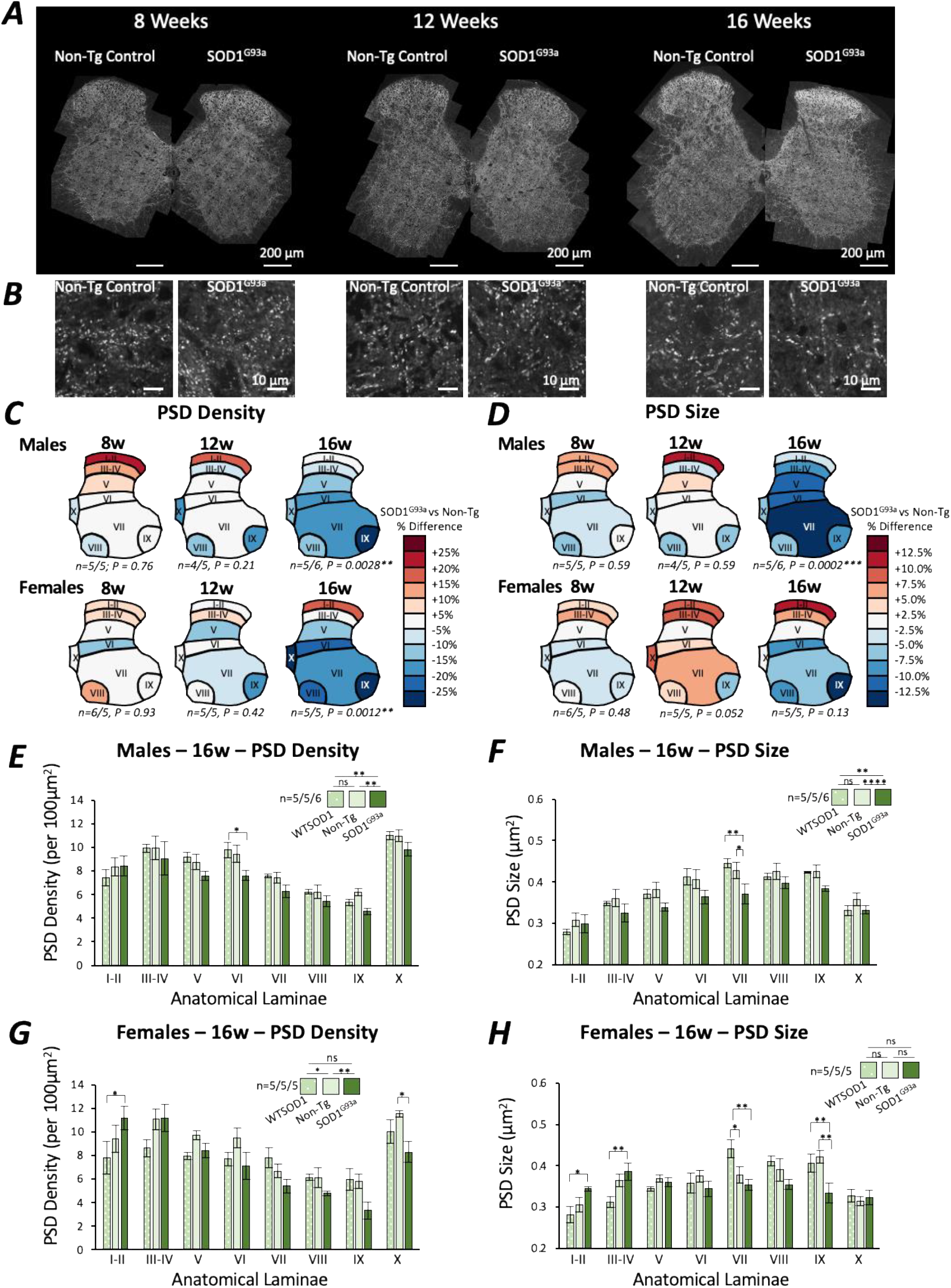
Mapping excitatory synapses in the spinal cord revealed early symptomatic stage synaptic changes in SOD1^G93a^ mice. **A**. High-resolution mapping of PSD95-eGFP was performed to analyse synaptic changes in SOD1^G93a^ mice at different time points of the disease. **B**. Example images of postsynaptic puncta in non-Tg control and SOD1^G93a^ mice at different time points of disease. **C**. Heat maps of PSD density in male and female SOD1^G93a^ mice compared to their respective non-Tg controls at different ages. Sample sizes denoted as N = number of non-Tg controls / number of SOD1G93a’s. **D**. Heat maps of PSD size in male and female SOD1^G93a^ mice compared to their respective non-Tg controls at different ages. **E**. Chart plotting PSD density in all 8 laminae of 16-week-old non-Tg controls, WTSOD1 controls and SOD1^G93a^ males. Sample sizes denoted as N = number of WTSOD1’s / number of non-Tg controls / number of SOD1^G93a^’s. **F**. Chart plotting PSD size in all 8 laminae of 16-week-old non-Tg controls, WTSOD1 controls and SOD1^G93a^ males. **G**. Chart plotting PSD density in all 8 laminae of 16-week-old non-Tg controls, WTSOD1 controls and SOD1^G93a^ females. **H**. Chart plotting PSD size in all 8 laminae of 16-week-old non-Tg controls, WTSOD1 controls and SOD1^G93a^ females.

The SOD1^G93a^ 16-week old females displayed reduced PSD density when compared to non-Tg control females (F(1, 64) = 10.26, P=0.0021), but no difference in PSD size (F (1, 64) = 3.232, P=0.077). Furthermore, these effects were not significant in comparison to the female 16-week-old WTSOD1 mice. Interestingly, it was noted that unlike the SOD1^G93a^ males, the SOD1^G93a^ females displayed more complex laminae-genotype-specific synaptic changes in both PSD density (F (14, 96) = 2.337, P=0.008) and size (F (14, 96) = 3.675, P<0.0001). The dorsal laminae I-II of SOD1^G93a^ females displaying increased PSD density and size compared to controls, while ventral laminae were more likely to show reduced PSD density and size compared to controls.

To address whether synaptic loss was associated with MN cell death, we performed immunolabelling using choline acetyltransferase (ChAT) specific antibody. We found no significant loss of MNs in 16-week old SOD1^G93a^ male spinal cords (t(6)=0.99, p=0.36). While other studies indicate MN cell loss can occur from approximately 16 weeks onwards (Feeney et al., 2001), our data suggest that synaptic changes occur prior to overt MN cell death (SI. Fig. 2A-D).

Next, we examined the spinal cords of mice carrying a mutation in the C9orf72 gene, one of the most common genes associated with familial and sporadic ALS (SI. Fig 3A-B). Synaptic maps from 12 week and 22 week old C9orf72 mutant mice of both sexes were obtained to capture reported pre-symptomatic and early-symptomatic stages of the disease phenotype in this model (Liu et al., 2016; Mordes et al., 2020; Nguyen et al., 2020). Mapping PSD95-eGFP synapses revealed no difference in PSD density or size in C9orf72 mice compared to non-Tg controls at either 12 or 22 weeks of age in both males and females (SI. Fig. 3C-J). The lack of synaptic changes in C9orf72 mice may reflect the fact that no motor phenotypes were observed in any of our C9orf72 mice even by 22 weeks. Our observations are consistent with reports from other groups that the penetrance of the phenotype in this mouse model can be very low in some colonies (Mordes et al., 2020; Nguyen et al., 2020).

In summary, our extensive synaptic mapping reveals a loss of PSDs at excitatory synapses in both male and female SOD1^G93a^ mice during early symptomatic stages. Our data also suggest that male SOD1^G93a^ mice display structural changes in synapses, an effect not as clearly observed in female SOD1^G93a^ mice, perhaps due to the delayed onset of disease in females. While we observed no synaptic changes in C9of72 mutant mice, this mirrors the fact that no ALS-related phenotypes, such as acute motor decline or weight loss, were observed in these mice. Given that our C9orf72 mice fail to robustly recapitulate an ALS disease phenotype, they were not included in any further analyses.

### Super-Resolution Microscopy Reveals Loss of Multi-Nanocluster Synapses in ALS Mouse Spinal Cord

Our high-resolution microscopy data revealed both a reduced number and reduced average size of PSDs in 16-week SOD1^G93a^ males. While confocal-level resolution is sufficient to measure synapse number and overall PSD size, the nanoscale organisation of synapses is best resolved using super-resolution microscopy. PSD95 forms nanoclusters (NCs) of approximately 100-150 nm diameter within the PSD (Broadhead et al., 2016, 2020; MacGillavry et al., 2013; Nair et al., 2013). PSD size, which correlates with synaptic strength, is determined by the number of NCs per PSD (Fig. 2A). We performed g-STED microscopy of PSD95-eGFP expression in 16-week-old male SOD1^G93a^ and non-Tg control mice to quantify synaptic nanostructure (Fig. 2B; n=4 mice per group). PSD95 NC size did not differ between genotypes (diameter ∼140nm; Fig. 2C; t(6)=0.31, p=0.77), and was consistent with previous measurements of PSD95 NC size using g-STED microscopy in both the brain and spinal cord (Broadhead et al., 2016, 2020; Fukata et al., 2013). Instead, SOD1^G93a^ mice showed a reduced average number of NCs per PSD (∼1.75 NCs per PSD) compared with non-Tg controls (∼2 NCs per PSD) (Fig. 2D; t(6)=2.8, p=0.03). This appears to be the result of a significant loss of large multi-NC-PSDs in SOD1^G93a^ mice, while smaller 1 or 2 NC-PSDs were observed with similar frequency (Fig. 2E; χ(2) = 14.7, p=0.0006). In non-Tg controls, 23% of synapses were classed as containing 3 or more NCs, while 17% of SOD1^G93a^ synapses contained 3 or more NCs.

**Fig. 2.**
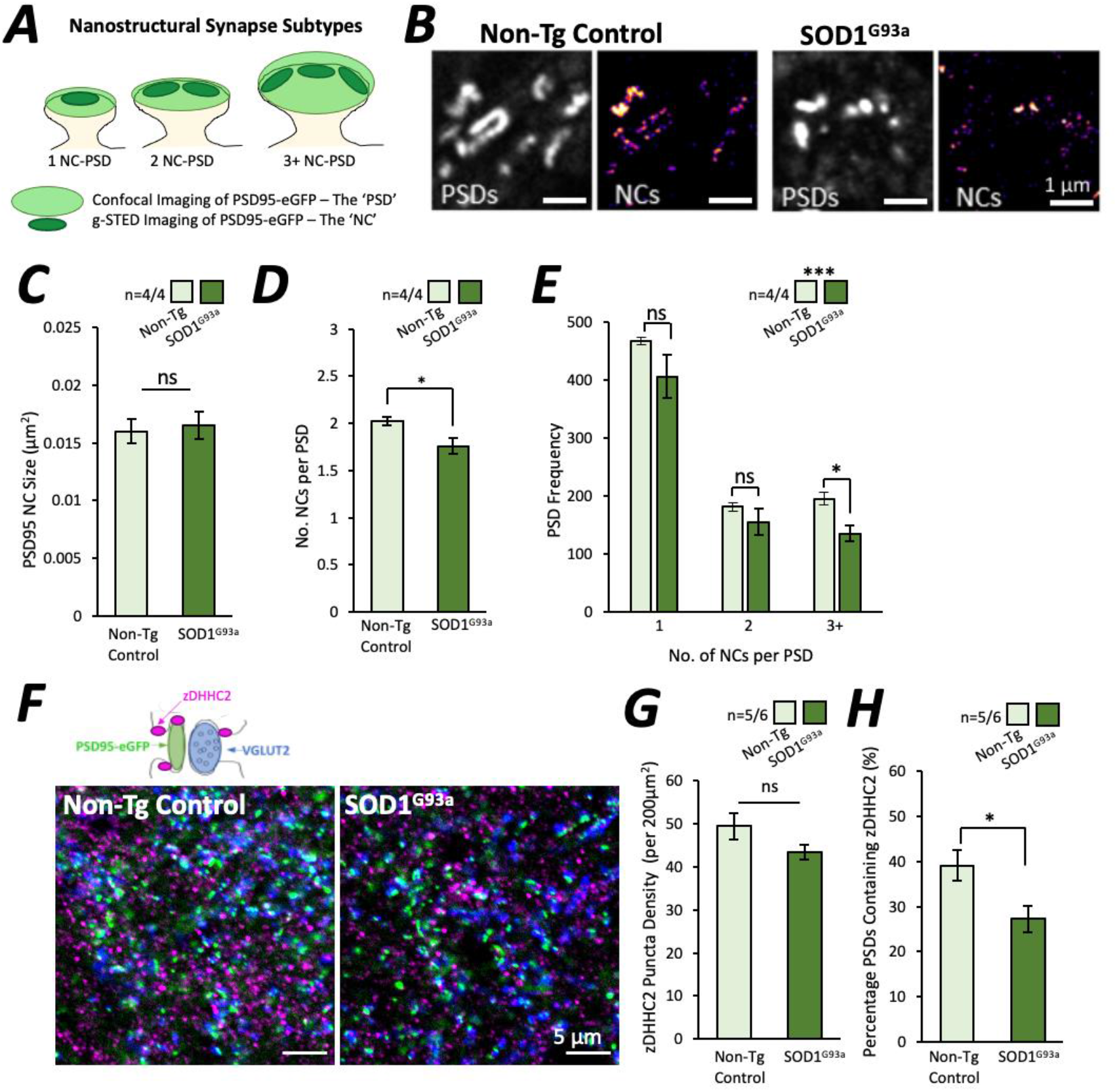
Nanoscopic Changes in Substructural Organisation of PSD95 in ALS Synapses. **A.** Diagram of synapse subtypes, defined by the number of PSD95 nanostructures (NCs) per PSD. PSDs can be visualised using confocal microscopy while NCs are resolved from correlative g-STED images. **B**. Spinal cord tissue from non-Tg control and SOD1^G93a^ mice were imaged using confocal and g-STED microscopy to visualise PSDs and NCs respectively. **C**. Graph showing the size of the PSD95 NCs was not different between non-Tg controls and SOD1^G93a^ mice. **D**. Graph showing the average number of NCs per PSD was significantly reduced in SOD1^G93a^ mice. **E**. Graph showing the number of large 3+ NC-PSDs are specifically reduced in SOD1^G93a^ mice compared to non-Tg controls. **F**. High-resolution imaging of synapses (PSD95 and VGLUT2) and the palmitoylation enzyme, zDHHC2, was performed to investigate whether changes in zDHHC2 expression might explain the reduced number of PSD95 NCs at synapses. **G**. Graph showing no difference in zDHHC2 puncta density between control and SOD1G93a mice. **H**. Graph showing reduced number of PSDs expressing zDHHC2 in SOD1^G93a^ mice.

The formation of PSD95 NCs at the synapse is dependent on palmitoylation of PSD95 at its N-terminal cysteine residue by the palmitoylating enzyme, zDHHC2. Having revealed a reduced number of PSD95 clusters per synapse in ALS mice, we assessed whether a change in zDHHC2 expression could be responsible. We analysed excitatory synapses (PSDs using PSD95-eGFP and presynaptic terminals using anti-VGLUT2 immunolabelling) for their expression of zDHHC2 by acquiring high-resolution images from the ventral horn of spinal cords from 16 week male non-Tg control and SOD1^G93a^ mice (Fig 2F). zDHHC2 displayed a punctate expression showing close association with both PSD95-eGFP and VGLUT2 puncta. While the overall density of zDHHC2 puncta was not significantly different between genotypes (Fig 2G; t(9)=2.1, p=0.067), there was a reduced number of PSDs colocalising with z-DHHC2 in SOD1^G93a^ mice (Fig. 2H; t(9)=2.6, p=0.031). This finding suggests that reduced postsynaptic presence of the palmitoylation enzyme, zDHHC2, may be related to a reduced number of PSD95 NCs at the synapse and lead to incremental reductions in postsynaptic strength.

### Widescale Mapping Reveals Tripartite Synapse Loss in ALS Mouse Spinal Cord

Tripartite synapses are chemical synapses contacted by perisynaptic astrocytic processes (PAPs). Previously we used antibody labelling against PAP proteins EAAT2 and p-Ezrin to identify tripartite synapses in the mouse lumbar spinal cord based on colocalization or close association of PAP proteins with postsynaptic PSD95 and presynaptic VGLUT2 (Broadhead et al., 2020). p-Ezrin and EAAT2 puncta strongly colocalised with one another in the mouse spinal cord (SI. Fig 4A). Both PAP markers are closely associated with astrocytic cell bodies, branches and fine processes, labelled with glutamine synthetase, but display little direct colocalization with astrocytic GFAP-positive primary arbours (SI. Fig 4B). These observations further confirm that both astrocytic EAAT2 and p-Ezrin label PAPs, making them ideal markers, in combination with synaptic markers, for visualising tripartite synapses. We, therefore, performed extensive high-resolution mapping of tripartite synapses in male SOD1^G93a^ and non-Tg control spinal cords, labelling PSDs (PSD95-eGFP), presynaptic terminals (VGLUT2) and PAPs (EAAT2 or p-Ezrin).

At 8 weeks, SOD1^G93a^ mice displayed an increased density of EAAT2 puncta (F (1, 56) = 11.19, p=0.0015; SI. Fig 5A) and p-Ezrin puncta (F (1, 56) = 4.672, p=0.035; SI. Fig. 5B) compared to non-Tg controls. No changes were observed in 12-week-old mice (SI. Fig 5C-D), but by 16 weeks, SOD1^G93a^ mice displayed significantly reduced EAAT2 (F (1, 64) = 11.31, p=0.0013; SI. Fig 5E) and p-Ezrin (F (1, 48) = 8.828, p=0.0046; SI. Fig. 5F) puncta density compared to non-Tg controls. These data demonstrate age-dependent changes in PAP proteins, suggesting an early stage increase in p-Ezrin-positive PAPs and increased EAAT2 expression at PAPs, followed by a significant loss of both proteins by 16 weeks, following the onset of early motor deficits.

We next quantified the density of non-tripartite synapses and tripartite synapses in non-Tg control and SOD1^G93a^ spinal cords at 8, 12 and 16 weeks. Using p-Ezrin, we observed no difference in the density of non-tripartite synapses between non-Tg controls and SOD1^G93a^ mice at either 8 weeks (Fig. 3C, F (1, 56) = 2.236, P=0.1404) 12 weeks (Figure 3E; F(1,56)=0.5631, P=0.456) or 16 weeks of age (Fig 3G; F(1,48)=2.4, p=0.126). The density of tripartite synapses was also no different between non-Tg controls and SOD1^G93a^ at 8 weeks (Fig 3C; F(1,56)=0.2998, P=0.586) and 12 weeks (Fig. 3E; F(1,56)=0.833, p=0.365). However, there was a significant reduction in tripartite synapse density at 16 weeks in SOD1^G93a^ mice (Fig 3G; F(1,48)=14.59, p=0.0004).

**Fig. 3.**
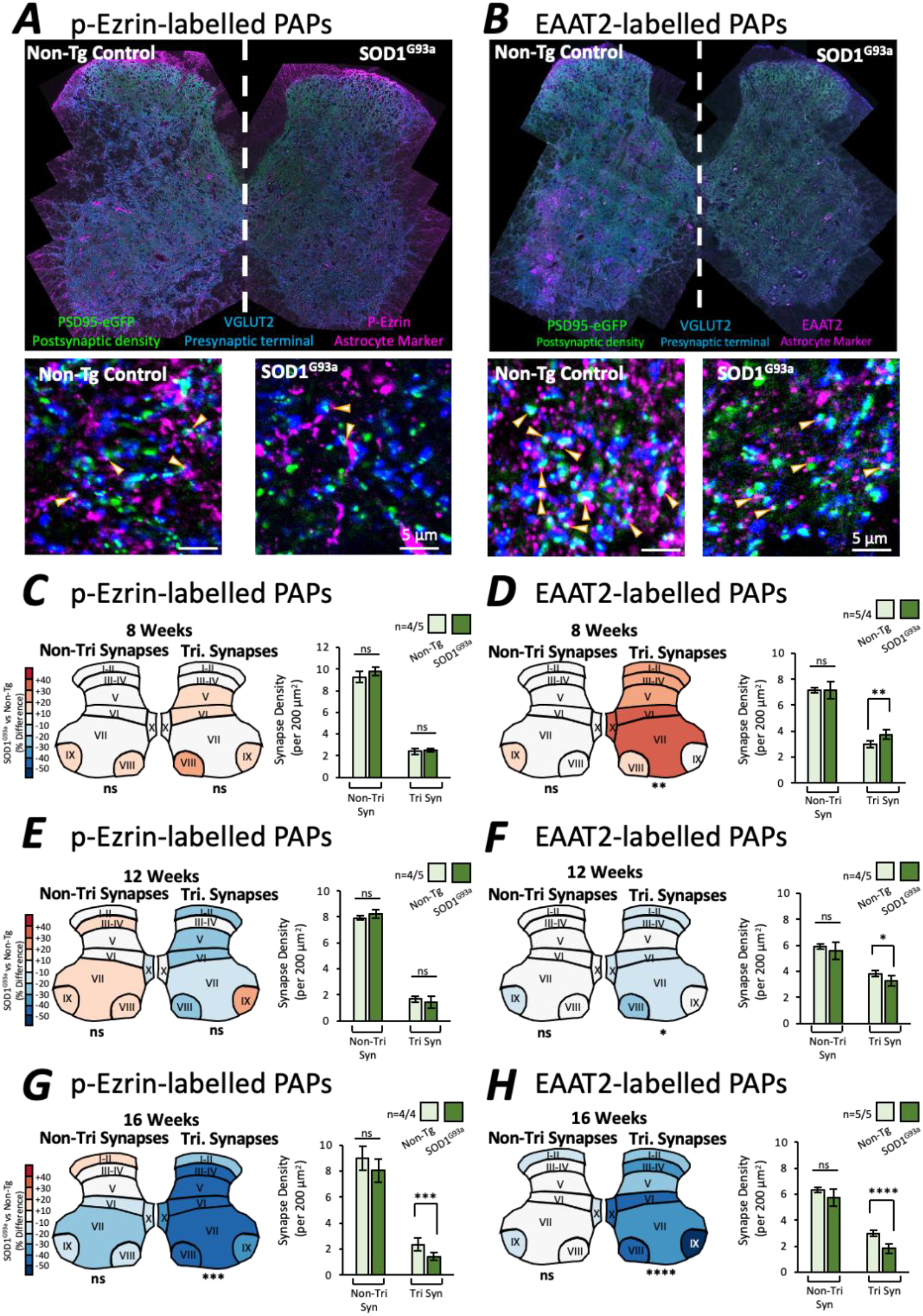
Mapping Tripartite Synapses in SOD1^G93a^ mouse Spinal Cord. **A.** Example of high-resolution spinal cord mapping of tripartite synapses from 16-week-old male non-Tg control and SOD1^G93a^ mice using PSD95-eGFP, VGLUT2 and p-Ezrin to label the PSD, presynaptic terminal and PAPs respectively. Cropped images display individual synaptic and tripartite synaptic structures. **B**. Example of high-resolution spinal cord mapping of tripartite synapses from 16-week-old male non-Tg control and SOD1^G93a^ mice using PSD95-eGFP, VGLUT2 and EAAT2 to label the PSD, presynaptic terminal and PAPs respectively. Cropped images display individual synaptic and tripartite synaptic structures. **C**. Colour coded heat maps denote the difference in synapse or tripartite synapse density observed in SOD1^G93a^ mice as a percentage of the respective non-Tg controls. Bar charts display the mean synapse density from mice, averaged across all anatomical subregions for simplicity. From p-Ezrin labelling, no changes in non-tripartite or tripartite synapses were observed in 8 week SOD1^G93a^ mice. **D**. From EAAT2 labelling, no changes in non-tripartite synapses were observed, but a significant increase in tripartite synapse numbers were observed in 8 week SOD1^G93a^ mice compared to control. **E**. From p-Ezrin labelling, no changes in non-tripartite or tripartite synapses were observed in 12 week SOD1^G93a^ mice. **F**. From EAAT2 labelling, no changes in non-tripartite synapses were observed, but a significant decrease in tripartite synapse numbers were observed in 12 week SOD1^G93a^ mice compared to control**. G.** From p-Ezrin labelling, no changes in non-tripartite synapses were observed, but a significant decrease in tripartite synapse numbers were observed in 16 week SOD1^G93a^ mice compared to control. **H.** From EAAT2 labelling, no changes in non-tripartite synapses were observed, but a significant decrease in tripartite synapse numbers were observed in 16 week SOD1^G93a^ mice compared to control

Similarly, using EAAT2 as the PAP marker, the density of non-tripartite synapses was no different between non-Tg controls and SOD1^G93a^ mice at either 8 weeks (Fig. 3D; F(1, 56)<0.0001, p=0.99), 12 weeks (Fig. 3F; F (1, 56) = 1.371, p=0.247) or 16 weeks (Fig. 3H; F(1,64)=2.403, p=0.126). The density of EAAT2-associated tripartite synapses, however, was significantly increased in 8 week SOD1^G93a^ mice (Fig. 3D, F (1, 56) = 7.214, p=0.0095), then subtly but significantly reduced in 12 week SOD1^G93a^ mice (Fig. 3F; F (1, 56) = 4.774, p=0.033) and more significantly reduced in 16 week SOD1^G93a^ mice (Fig. 3H; F (1, 64) = 23.58, p<0.0001). From these experiments, using two different PAP markers, our data consistently demonstrate that tripartite synapses are a selectively vulnerable synapse subtype in early-symptomatic stage ALS mice.

### Selective loss of Tripartite Synapses in Human ALS Spinal Cord

Finally, we addressed whether human cases of ALS also demonstrated synaptic and tripartite synaptic degeneration. Post-mortem cervical spinal cord sections were obtained from human cases, including 6 healthy controls (5 male, 1 female), and 9 ALS patients (all male) comprising 4 SOD1 cases and 5 C9orf72 cases (SI. Table 1). Synapses and tripartite synapses were visualised in the ventral horn using immunolabelling for PSD95 and p-Ezrin to identify PSDs and PAPs respectively (Fig. 4A-B).

**Fig. 4.**
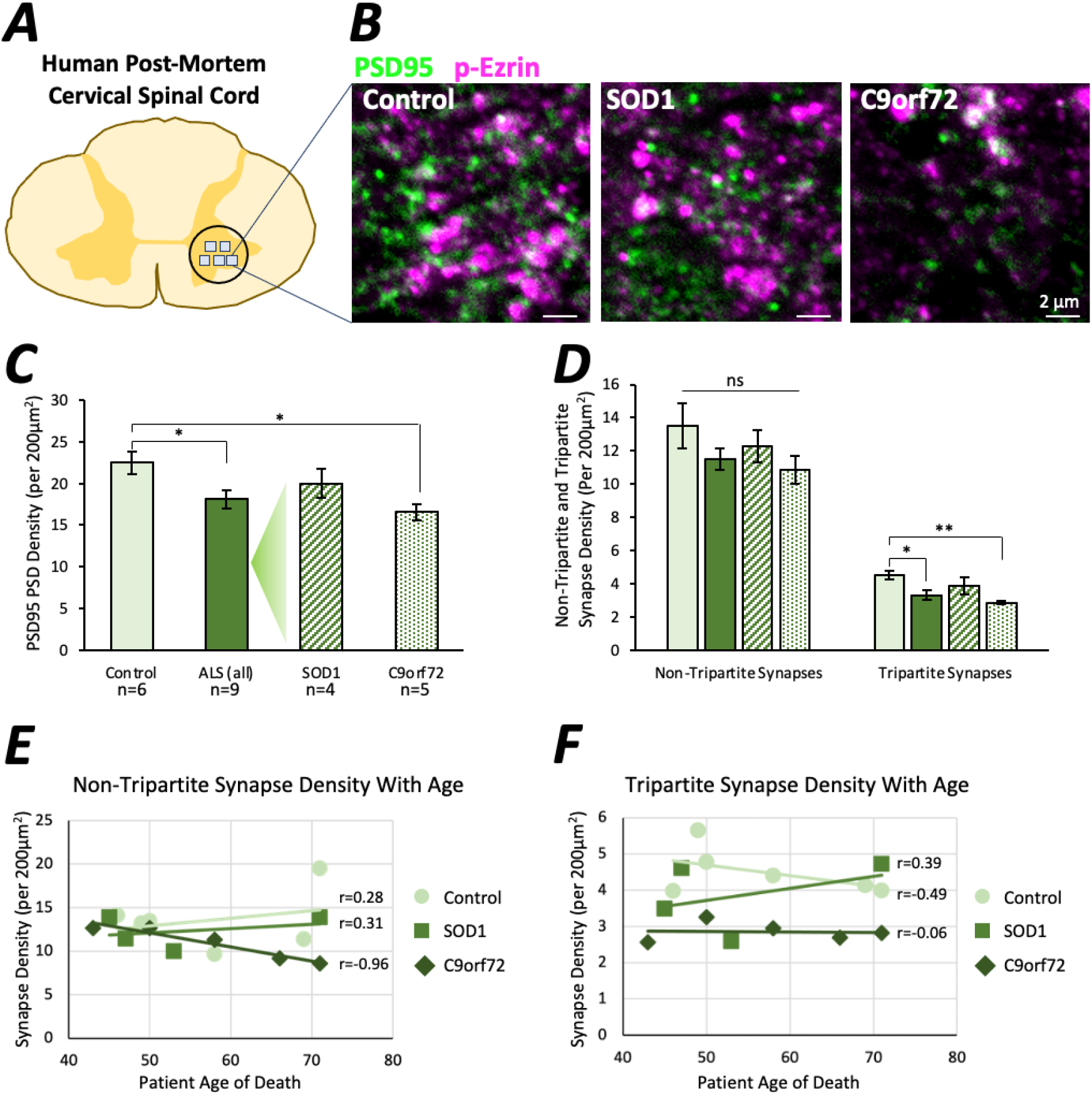
Tripartite Synapses are selectively lost in the cervical spinal cord of human ALS cases. **A.** Diagram of human cervical spinal cord and location of image sampling. **B**. Example images of synapses (PSD95 – green) and astrocytic processes (p-Ezrin – magenta) in human spinal cord tissue. **C**. Graph showing the total synapse density (all PSD95 puncta) in controls, all ALS cases, and then SOD1 and C9orf72 cases separated. **D**. Graph showing synapse density in patient groups when synapses are separated based on whether they are non-tripartite synapses or tripartite synapses, as determined by the association of PSD95 with p-Ezrin. **E**. Graph showing the correlation between individual patient ages and non-tripartite synapse density for patient subgroups. **F**. Graph showing the correlation between individual patient ages and tripartite synapse density for patient subgroups.

When comparing control cases with all ALS cases (SOD1 and C9orf72), PSD density was significantly reduced in the ALS cervical spinal cord (Fig. 4C; t(13)=2.6, p=0.021). Interestingly, when control, SOD1 and C9orf72 cases were analysed separately, only C9orf72 cases displayed significantly reduced PSD density compared to controls (F(2,12)=5.4, P=0.0214; Control versus C9orf72: p=0.0167). We observed no difference in PSD size between the three groups (F(2,12)=0.16, P=0.85).

Next, we asked whether tripartite synapses were affected in ALS patients. Similar to results from the SOD1^G93a^ mouse model, the density of non-tripartite synapses was no different between control and ALS patients (F(2,12)=1.44, P=0.275), while the density of tripartite synapses was significantly reduced in C9orf72 cases compared to controls (F(2,12)=8.163, P=0.0058, Control versus C9orf72: p=0.0044).

When combining all patient cases, synapse density measurements showed no correlation with patient age (total synapse density: R^2^ = 0.003, p=0.855; non-tripartite synapse density: R^2^ = 0.002, p=0.887; tripartite synapse density: R^2^ = 0.002, p=0.867). Examining C9orf72 patient cases only, there appeared to be a negative correlation between age and the density of synapses (R^2^=0.769, p=0.051) and non-tripartite synapses (R^2^=0.926, p=0.0087) (Fig. 4E). As a result, total synapse density was more notably reduced in older C9orf72 patients compared to age-matched control cases, while total synapse density in younger C9orf72 patients was more similar to age matched controls. In contrast, the density of tripartite synapses was not correlated with age (R^2^=0.004, p=0.920), and remained lower than control cases across all ages (Fig 4F).

Taken together these findings provide evidence that tripartite synapses are selectively vulnerable to degeneration in human ALS cases, as observed in the SOD1^G93a^ mouse model. Furthermore, tripartite synapse loss, as opposed to merely synapse loss, may be a more reliable biomarker of ALS, as synapse density could be confounded by age-related changes in other non-tripartite synapses.

## Discussion

This study set out to investigate how synapses and tripartite synapses are affected in the ALS spinal cord. To this end, we performed large-scale quantitative mapping of synapses and tripartite synapses in ALS mouse models and human post-mortem tissue. Our data revealed that, of the millions of synapses in the spinal cord, tripartite synapses appear to be selectively vulnerable to degeneration in ALS. We therefore describe ALS as a tripartite synaptopathy.

The functional, structural and molecular diversity of synapses throughout the nervous system, i.e. synaptome diversity, endows the nervous system with the capability to support a wide repertoire of behaviours from motor control to learning and memory (Grant, 2019; Grant & Fransén, 2020). Over 130 different nervous system disorders, including ALS, can arise from mutations in postsynaptic proteins (Bayés et al., 2011), and lead to wide-scale reprogramming of the synaptome (Zhu et al., 2018). Different diseases of the nervous system may display a unique synaptome signature, as certain subtypes of synapses may be more vulnerable to changes in one condition compared to another – thus leading to the specific behavioural consequences and symptoms of a given condition. By first identifying the vulnerable synapse subtypes of a given disease, we can establish therapeutic roadmaps to selectively target these synapse subtypes to treat the symptoms of disease.

The PSD95-eGFP mouse model provides a powerful tool with which to investigate the effects of diseases on synapses as PSD95 is abundantly expressed at the PSDs of excitatory synapses and forms a core molecular framework for synaptic machinery (Cizeron et al., 2020; Fernández et al., 2009; Frank & Grant, 2017; Husi & Grant, 2001; Zhu et al., 2018). The same mouse model can be used for nanoscopic analyses of synapse subtypes based on postsynaptic substructure using super-resolution microscopy (Broadhead et al., 2016, 2020).

This study also sought to identify whether astrocytes play a role in the vulnerability of synapses in ALS. Labelling the astrocytic component of the tripartite synapse was performed here using immunolabelling for either EAAT2 or p-Ezrin. Various studies support the use of either EAAT2 or Ezrin/p-Ezrin as robust markers for the PAPs which contact synapses (Derouiche et al., 2002; Foster et al., 2018; Heller & Rusakov, 2017; Lavialle et al., 2011). By performing tripartite synapse mapping with both PAP markers in parallel, we have observed strikingly similar findings which give us considerable confidence in our conclusion that tripartite synapses are vulnerable in ALS. Reduced levels of EAAT2 in the ALS spinal cord have been well documented (Howland et al., 2002; Lin et al., 1998; Sasaki et al., 2000). The consistent reduction in both EAAT2 and the actin filament binding protein, p-Ezrin, suggest not just a molecular change in astrocytic protein expression but that the PAPs themselves are being lost.

Using the SOD1^G93a^ mouse model, we observed synapse loss, structural changes in PSDs, changes in PAPs and a loss of tripartite synapses at early, symptomatic stages. While we observed significant synaptopathy in SOD1^G93a^ mouse spinal cords, we observed no such alterations in C9orf72 mutant mice. The C9orf72 (C9BAC-500) mice were originally reported to display molecular hallmarks of ALS pathology, and acute motor deficits in 35% of the females carrying the mutation (Liu et al., 2016). Further investigation by other groups using this model have shown different degrees of phenotypic penetrance (Mordes et al., 2020; Nguyen et al., 2020). In our hands, the C9orf72 mice (the pure-bred colony and the progeny of PSD95-eGFP cross-breeding) display no such acute motor phenotypes in either males or females, though we did observe RNA foci accumulating in cell nuclei – a key molecular hallmark of ALS pathology. We consider that either genetic or environmental differences in our experimental colonies may have led to significantly reduced penetrance of the disease phenotype in the mouse line (Nguyen et al., 2020). Given that phenotypic penetrance can be low, and that acute ALS phenotypes can occur between 20 and 40 weeks of age, further data collection from larger numbers of older animals may have captured the potential synaptic phenotype of this model. On the other hand, another mouse model with a more reliable phenotype, such as TDP43^A315T^ which has been shown previously to display both synaptic and astrocytic changes (Jiang et al., 2019; Ke et al., 2015), may be a more suitable alternative for further investigation of tripartite synaptopathy.

We employed large-scale anatomical mapping of synapses and tripartite synapses in the lumbar mouse spinal cord. It was anticipated that some anatomical laminae, for example laminae VII-IX where MNs reside, may display more significant changes in synapses in ALS. Despite significant inter-regional diversity in PSD size and number (Broadhead et al., 2020), our data suggest that the loss of synapses occurs widely throughout the spinal cord and not simply localised to regions where MNs reside. This observation suggests that ALS may impact a broad range of sensory, integratory and motor circuitry. Interestingly, female SOD1^G93a^ mice displayed significantly increased numbers of synapses in dorsal laminae I-II compared to controls, which is supported by previous reports of sensory-level alterations in ALS and other related disorders (Fletcher et al., 2017; Vaughan et al., 2018). Our mapping analyses also suggested that while PSD loss occurred in both male and female SOD1^G93a^ mice, structural changes in PSDs may be more prominent in males. Evidence from other studies also indicates a gender bias in SOD1^G93a^ mice, with males showing a slightly earlier onset and increased rate of disease progression, and some changes in specific subtypes of spinal synapses only occurring in males (Cacabelos et al., 2016; Heiman-Patterson et al., 2005; Herron & Miles, 2012). In addition, clinical ALS cases present with a higher incidence in males (McCombe & Henderson, 2010).

Our extensive analysis of SOD1^G93a^ mouse spinal cords revealed significant postsynaptic changes at the early symptomatic stage of 16 weeks (112 days), with no overt changes at 12 weeks (84 days) or 8 weeks (56 days) of age, although early signs of tripartite synapse loss were observed at 12 weeks. Others have reported reduced sizes of clusters of other synaptic proteins such as scaffolding proteins Shank1 and Homer, and glutamatergic AMPA receptors on spinal cord MNs in pre-symptomatic 8 week old SOD1^G93a^ mice (Bączyk et al., 2020). Similarly, pre-symptomatic changes in cortical synapses and neuromuscular junctions have been observed as early as 4-6 weeks in mouse models (Eisen, 2021; Fischer et al., 2004; Fogarty et al., 2015, 2016; Frey et al., 2000). Our current data does not suggest that significant synaptic alterations occur pre-symptomatically in spinal cord circuits, but instead suggests that they occur at the early-symptomatic stages of the disease. Dying-forward (cortex to spinal cord) and dying-backward (periphery to spinal cord) mechanisms exist pre-symptomatically (Eisen, 2021; Fischer et al., 2004), and may not lead to significant synaptic changes in the spinal cord until later in the disease progression. Furthermore, different techniques to visualise and quantify synaptic structure and function may confound direct comparisons across studies. For example, it is conceivable that changes in dendritic spines may occur without any apparent changes in the expression or organisation of PSD95 (Fogarty et al., 2015). Comparative analyses of synaptic changes in both the brain and spinal cord in the early stages of ALS may help decipher whether synaptic changes occur simultaneously in various parts of the CNS, or whether, for example, motor cortex dysfunction drives synaptic dysfunction in the spinal cord in a feedforward manner.

In accordance with other reports of changes in the structure of synapses in ALS (Bączyk et al., 2020; Sasaki & Iwata, 1995, 1996), our high-resolution imaging revealed reduced PSD size in early-symptomatic SOD1^G93a^ males. By employing super-resolution microscopy, we have performed nanoscopic analysis of the molecular organisation of synapses in ALS for the first time. This powerful microscopy technique has characterised the nature of ALS synaptopathy at the sub-diffraction scale, revealing a reduced number of large multi-NC synapses. PSD95 forms nanoscale postsynaptic domains that tether neurotransmitter subdomains at the postsynaptic membrane, which align with presynaptic release sites to create trans-synaptic signalling nanocolumns (Tang et al., 2016). Synaptic strength appears to correlate with the number of nanodomains within the synapse (Broadhead et al., 2016; MacGillavry et al., 2013; Nair et al., 2013). Our previous work has shown that multi-NC synapses are more likely to be tripartite synapses, based on their association with PAP proteins EAAT2 and p-Ezrin (Broadhead et al., 2020). This supports the concept that large multi-NC tripartite synapses are those which are selectively vulnerable to degeneration in ALS. Given that the number of PSD95 subdomains per synapse correlates with synaptic strength, it is possible that the loss of these large, high-fidelity synaptic connections significantly contributes to early-stage, progressive motor deficits. zDHHC2 is a palmitoylating enzyme that facilitates the clustering of PSD95 at the synapse. From our data, we also speculate that the dissociation of zDHHC2 from the PSD may lead to reduced clustering of PSD95 at the synapse (Fukata et al., 2013), though the precise role of protein palmitoylation in neurodegenerative disease remains to be determined (Cho & Park, 2016; Zaręba-Kozioł et al., 2018).

Importantly, we were able to replicate our key observations of synapse and tripartite synapse loss in ALS mouse models when studying human cervical spinal cord tissue from ALS patients. Our analysis suggested that C9orf72 patients displayed significant synaptic loss and tripartite synapse vulnerability, while SOD1 patients did not show significant synapse or tripartite synapse loss compared to controls. ALS is a clinically and molecularly heterogenous disorder, and C9orf72 mutations represent the most common genetic basis of both sporadic and familial ALS. A previous investigation into synaptic changes in the prefrontal cortex of ALS patients revealed no differences in synaptic loss between SOD1 and C9orf72 associated cases, but did suggest that synaptic loss was most significant in cases displaying TAR-DNA Binding protein (TDP43) pathology – of which SOD1 cases showed none (Henstridge et al., 2018).

The mechanisms leading to tripartite synapse degeneration remain to be determined. Mislocalisation and aggregation of TDP43 in the cytoplasm of both neurons and glia is correlated with synapse loss in human ALS (Henstridge et al., 2018), and could play a role in designating tripartite synapses for degeneration, though there is mixed evidence of TDP43 pathology in the SOD1^G93a^ mouse model (Jeon et al., 2019; Shan et al., 2009; Tan et al., 2007). Alternatively, the loss of astrocytic EAAT2 in ALS, due to aberrant RNA processing, is known to induce glutamatergic excitotoxicity that could lead to synapse degeneration and ultimately cell death (Howland et al., 2002; Lin et al., 1998; Maragakis & Rothstein, 2006; Rothstein et al., 1996). Our own data supports this hypothesis given that synaptic loss coincides with EAAT2 loss. Astrocytes also contribute to bi-directional signalling mechanisms that help modulate spinal cord mediated motor behaviour through purinergic signalling (Acton & Miles, 2015; Broadhead & Miles, 2020, 2021; Carlsen & Perrier, 2014; Witts et al., 2012, 2015). Neuronal-derived glutamate release during locomotor network activity activates astrocytes through mGluR5 receptors, leading to ATP/Adenosine release from astrocytes that inhibits neuronal activity through A1 adenosine receptors (Broadhead & Miles, 2020). Increased Adenosine levels in ALS patients (Yoshida et al., 1999), combined with increased expression of the excitatory A_2A_ Adenosine receptors (Ng et al., 2015), could imbalance this modulatory mechanism, biasing the system toward excitotoxic actions at the tripartite synapse. Our current study provides evidence that tripartite synapses, and the specific signalling pathways they utilise to regulate neuronal activity, may act as a fulcrum of ALS. These novel findings regarding selective synaptic vulnerability highlight numerous future avenues to explore and provide the framework to investigate the mechanisms and therapeutic potential of ALS tripartite synaptopathy.

## Acknowledgements

We would like to acknowledge the following funders: Motor Neurone Disease (MND) Association UK (Miles/Apr18/863-791), the Euan MacDonald Centre and Chief Scientist Office, The European Research Council (ERC) under the European Union’s Horizon 2020 Research and Innovation Programme (695568 SYNNOVATE), Simons Foundation Autism Research Initiative (529085), and the Wellcome Trust (Technology Development grant 202932). We would also like to acknowledge the MRC Edinburgh Brain Bank for the provision of human post-mortem tissue.

## Data Availability

The data used in this manuscript is freely available upon request.

## Methods

### Animals and Ethics

All procedures performed on animals were conducted in accordance with the UK Animals (Scientific Procedures) Act 1986 and were approved by the University of St Andrews Animal Welfare and Ethics Committee. The following mouse lines were kindly provided by Dr Richard Mead, University of Sheffield: B6SJL-TgN(SOD1-G93A)1Gur/J (SOD1^G93a^), B6SJL-Tg(SOD1)2Gur/J (WTSOD1) and FVB/NJ-Tg(C9ORF72)500Lpwr/J (C9orf72). SOD1^G93a^ and WTSOD1 were maintained on a congenic C57bl/6 background strain, while C9orf72 mice were maintained on an FVB background mouse strain. The ALS mouse lines were crossed with PSD95-eGFP^+/+^ mice, originally obtained from Prof. Seth Grant (University of Edinburgh), to produce transgenic (Tg) ALS and non-Tg control offspring expressing PSD95-eGFP^+/-^. Unlike the other mouse lines, C9orf72 mice were produced on an FVB background. The progeny of C9orf72 x PSD95-eGFP mice were a first generation FVB/C57bl/6 cross, displaying a light brown fur coat and black eyes (indicating they did not possess the recessive retinal degeneration 1 allele of Pde6brd1 which causes blindness in FVB mice).

Disease phenotype was monitored by routine weighing and behavioural scoring from the age of 30 days old (P30), with more frequent checks performed once the mice were over P60 or when mice were showing disease phenotypes. Mice were scored 0-4 in severity; 4 showing no phenotype, 3 showing hind-limb splay and tremors upon tail raising, 2 showing gait abnormalities, 1 showing hind limb dragging, 0 showing inability to self-right after 30 seconds when placed on its back. Mice were euthanised through either cervical dislocation or perfusion if displaying a score of 2 or lower.

Overall, SOD1^G93a^ mice (including pure SOD1^G93a^ mice and PSD95-eGFP x SOD1^G93a^ mice) presented hind limb tremor and reduced hind-limb splay by 75.9 ± 5.3 days (16 mice). Male SOD1^G93a^ mice presented with a slightly earlier onset of hind limb tremor (73.6 ± 5.6 days) compared to females (78.2 ± 4.1 days) (t(16)=2.01, p=0.061). We observed no difference in the age of symptom onset between SOD1^G93a^ mice and the cross-bred PSD95-eGFP mice, nor have we observed any difference in the age of symptom onset within our colonies over time. Our data is remarkably consistent with those of the source colonies (Mead et al., 2011) and suggests that little-to-no reduction in transgene copy number had occurred within our breeding colonies or experimental cohorts. No such overt phenotypes were observed in either the pure or PSD95-eGFP bred WTSOD1 and C9orf72 mouse colonies.

### Mouse Tissue Collection

Mice were anaesthetised with pentobarbitol (30mg/kg dose; Dolethal), and the chest cavity opened to reveal the heart. The right atrium was severed and 10 ml ice cold 1x phosphate buffered saline (PBS) was perfused through the left ventricle, followed by 10ml 4% paraformaldehyde (PFA; Alfa Aesar). The lumbar spinal cord was then dissected and incubated for a further 3-4 hrs in 4% PFA before being incubated in sucrose 30% w/v for 24 to 72 hours at 4°C until sunk. Tissue was then cryo-embedded in OCT compound and stored at −80°C. Cryosections were obtained using a Leica CM1860 cryostat at 20 μm thickness and adhered to Superfrost Gold Plus glass slides (VWR).

### Mouse Tissue Processing and Immunohistochemistry

For visualisation of PSD95-eGFP only, slides were thawed at room temperature, briefly washed in PBS then deionised (DI) water before being dried and mounted in lab-made Mowiol or Prolong Glass anti-fade for super-resolution microscopy with a 1.5 thickness coverslip. For immunohistochemistry, sections were first heated at 37°C for 30 mins to aid the adherence of the tissue to the glass slides and reduce tissue loss during subsequent wash steps. Slides were washed 3 times in PBS before being blocked and permeabilised in PBS containing 3% Bovine Serum Albumin (BSA) and 0.2% Triton X100 for 2 hours at room temperature. Primary antibody incubation was performed in PBS containing 1.5% BSA, 0.1% Triton X100 for 2 nights at 4°C. Once incubated, slides are washed 5 times in PBS over the course of 1-2 hours. Secondary antibody incubation was performed in 0.1% Triton X100 for 2 hours at room temperature followed by a further 5 washes in PBS over the course of 1-2 hours. Dye labelling of astrocyte-specific primary antibodies (GFAP, EAAT2 and p-Ezrin) was performed with 10µg of each antibody using the Lightning-Link conjugation kits (Abcam) to address whether some proteins colocalised in astrocytes when the primary antibodies were raised in the same species (rabbit). See Table 1 for details on antibodies for mouse tissue experiments.

**Table 1.**
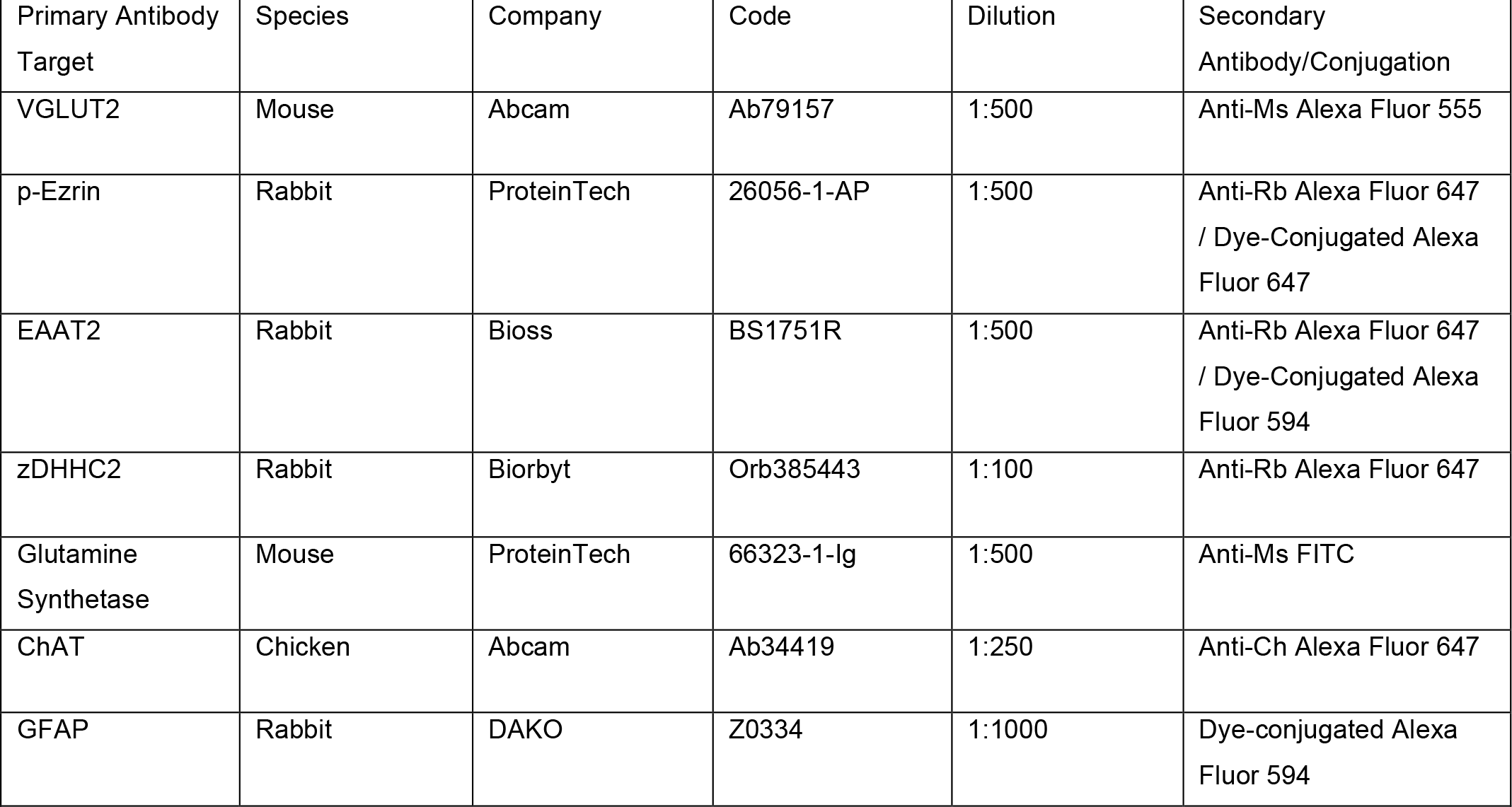
Mouse Spinal Cord Tissue Labelling

| Table 1. Mouse Spinal Cord Tissue Labelling |  |  |  |  |  |
| --- | --- | --- | --- | --- | --- |
| Primary Antibody Target | Species | Company | Code | Dilution | Secondary Antibody/Conjugation |
| VGLUT2 | Mouse | Abcam | Ab79157 | 1:500 | Anti-Ms Alexa Fluor 555 |
| p-Ezrin | Rabbit | ProteinTech | 26056-1-AP | 1:500 | Anti-Rb Alexa Fluor 647 / Dye-Conjugated Alexa Fluor 647 |
| EAAT2 | Rabbit | Bioss | BS1751R | 1:500 | Anti-Rb Alexa Fluor 647 / Dye-Conjugated Alexa Fluor 594 |
| zDHHC2 | Rabbit | Biorbyt | Orb385443 | 1:100 | Anti-Rb Alexa Fluor 647 |
| Glutamine Synthetase | Mouse | ProteinTech | 66323-1-Ig | 1:500 | Anti-Ms FITC |
| ChAT | Chicken | Abcam | Ab34419 | 1:250 | Anti-Ch Alexa Fluor 647 |
| GFAP | Rabbit | DAKO | Z0334 | 1:1000 | Dye-conjugated Alexa Fluor 594 |

### Human Post-Mortem Tissue and Immunohistochemistry

Human cervical spinal cord samples were obtained from the Edinburgh Brain and Tissue Bank, by Prof. Colin Smith. Samples of the cervical spinal cord were obtained from patients who died from ALS displaying mutations in the C9orf72 gene or the SOD1 gene. Age matched patients who died from natural causes were used as controls. Mean average patient ages were 57.2 years (controls), 57.6 years (C9orf72) and 54 years (SOD1). Sections were fixed, paraffin embedded and sectioned at 4 μm thickness and mounted to glass slides. For immunohistochemistry, samples were first de-paraffinised with sequential 3 minute washes in Xylene, 100% ethanol, 95% ethanol, 70% ethanol and 50% ethanol. Antigen retrieval was performed by incubating samples in Citric Acid (pH 6.0) for 20 minutes in a steamer, providing a consistent heat of 95°C. Following antigen retrieval, sections were washed 3 times in PBS and then incubated in 1x Tris buffered Saline (TBS) with 5% BSA, 0.2% Triton X100 for 2 hours at room temperature. Samples were then incubated in primary antibody solution of TBS with 3% BSA, 0.2% Triton X100 for 2 nights at 4°C before being washed 5 times in TBS containing 0.2% Triton X100 over the course of 1 hour. Secondary antibody incubation was performed in TBS with 0.2% Triton X100 solution for 3 hours at room temperature followed by 5-6 washes in PBS with 0.2% Triton X100 over the course of 2 hours. Finally, sections were washed in DI water, dried and mounted in Mowiol. See Table 2 for antibodies used in human tissue experiments.

**Table 2.** Human Spinal Cord Tissue Labelling

| Table 2. Human Spinal Cord Tissue Labelling |  |  |  |  |  |
| --- | --- | --- | --- | --- | --- |
| Primary Antibody Target | Species | Company | Code | Dilution | Secondary Antibody/Conjugation |
| PSD95 | Guinea Pig | Synaptic Systems | 124-014 | 1:500 | Gt Anti-Gp 647 |
| p-Ezrin | Rabbit | ProteinTech | 26056-1-AP | 1:250 | Gt Anti-Rb Cy3 |

### Primary Spinal Cord Astrocyte Cultures

Spinal cords were isolated from postnatal day (P) 2-4 mouse pups (C9orf72^+/-^ and non-Tg littermates X PSD95-eGFP^+/+^). Pups were euthanized by cervical dislocation, followed by decapitation and evisceration. Spinal cord isolation was performed using contents of ‘NeuroCult™ Enzymatic Dissociation Kit for Adult CNS Tissue (StemCell Technologies)’. Spinal cords were isolated, trimmed to the lumbar region and sliced into 1mm sized pieces in ice cold ‘Tissue Collection Media (TCS)’. Astrocyte media needed for maintenance and dissociation washes contained: Low glucose DMEM + 10% FBS + 1% antibiotic-antimycotic (Gibco). For enzymatic dissociation, 1.5ml of ‘Dissociation Enzyme’ was added to the tissue and incubated at 37 ℃ for 3 minutes with agitation for 15 second intervals every minute. After 3 minutes, ‘Inhibition Solution’ was added directly to the enzyme followed by 3x washes with Astrocyte media. Tissue was resuspended in Astrocyte media, gently triturated and left to settle. Single cells were collected from solution once settled. Repeated trituration, settling and collection steps were performed until all tissue was dissociated. Cells were plated into T75 or T25 flasks coated in Poly-D-Lysine based on the number of cords isolated. The next morning flasks were washed to remove debris, then fresh media was added. Flasks were fed every 2-3 days until the day of passage, and after day of passage fed every 4-5 days. Flasks were passaged when confluent using TrypLE (Gibco) and plated at a density of 100,000 cells per 13mm coverslip.

### RNA FISH

Cy5-labelled DNA probes were used against the G4C2 RNA repeat expansion. All solutions were made up in DEPC-treated water or PBS. Fixed spinal cord tissue was permeabilised with 0.1% Triton X100 for 1 hour. Tissue was treated with 2x saline sodium citrate (SSC) and 10% formamide in PBS for 5 minutes. Slides were then incubated with hybridization solution containing probes in 10 % formamide overnight at 37°C. After hybridization slides were washed once in 2x SSC, 10% formamide, for 30 mins at 37°C, then washed in 2x SSC for 5 mins at room temperature. DAPI was used at 1:15,000 in PBS. Primary cultured astrocytes were treated using the same protocol as above, but with only 10 mins of initial permeabilization with Triton X100. Images were captured on a Zeiss Airyscan LSM800 at 63x.

### High-Resolution Microscopy

High-resolution confocal-like microscopy was performed using a Zeiss Axio Imager M2 Microscope equipped with an Apotome.2.0, which enables an XY resolution of 320 nm (Broadhead et al., 2020). Illumination was provided by a HXP120 lamp, and images acquired using a digital MRm camera. Exposure times and illumination intensity were kept consistent for analysis within batches of data sets. To ‘map’ entire hemi-sections of spinal cords to study anatomical diversity between laminae, single optical sections were captured and tiled across half a transverse section of the spinal cord. Maps were captured using a single Z-stack plane approximately 3-5 µm depth into the tissue that was kept at the same depth throughout the entire map. Images were stitched together to create whole montage images that were subsequently analysed.

### Super-Resolution Microscopy

Confocal and gated-stimulated emission depletion (g-STED) microscopy was performed using a Leica SP8 SMD g-STED microscope available at the Edinburgh Super-Resolution Imaging Consortium hosted by Heriot Watt University. Excitation was provided by a CW super-continuum white light laser source at 488nm to excite eGFP with depletion provided by a 594 nm and 775 nm laser. Images were acquired with a 100x 1.4NA STED objective lens with an optical zoom set to provide optimal xy-resolution.

### Image Analysis

Image analysis was performed in FIJI (Fiji is just ImageJ) (Schindelin et al., 2012). High-resolution image analysis was performed as described previously (Broadhead et al., 2020). A standard mouse spinal cord anatomical atlas was used to guide the delineation of Rexed’s laminae (Watson et al., 2009). For clarity, our delineation of lamina VIII represents a heterogenous population of ventral neurons including medial MN pools, whilst laminae IX represents lateral MN pools. For mapping data sets, whole images were processed using a background subtraction and gaussian smoothing. For the PSD95-eGFP-only data sets of excitatory synapse mapping, automated Moments-based thresholding was applied. For data sets involving detection of immunolabelled markers of synaptic or astrocytic structures, manual thresholding was typically required, and the researcher was blinded to the genotype of the mouse to avoid unconscious bias. Minimum and maximum size thresholds were set to minimise false positive detections of small dim structures of a few pixels in size and large clusters that were likely to be lipofuscin aggregates. Colocalisation between structures in high-resolution data sets was determined from binarized images with the criteria of showing overlap by at least 1 pixel.

Nanostructural analysis of PSD95 organisation from g-STED imaging was analysed as described previously (Broadhead et al., 2016, 2020). Briefly, confocal and g-STED images are processed with background subtraction and gaussian smoothing. Structures resolved in g-STED were deemed to be PSD95 nanoclusters (NCs) while larger diffraction-limited structures resolved with confocal were deemed to be whole PSDs. Structural parameters such as size, shape and intensity were obtained for PSDs and NCs, as well as how many NCs resided within a given PSD.

### Statistical Analysis

Data analysis and graph preparation was performed in Microsoft Excel. Statistical analyses including Shapiro Wilks normality tests, Students T-test, Single and Multi-factorial (Two-Way) ANOVA, Pearson’s correlations were performed using either SPSS (IBM) or Prism (Graphpad). Sulak’s post-hoc comparison test was performed for Two-Way ANOVA’s where more than two groups were compared, while Tukey’s post-hoc comparison was performed in Two-Way ANOVA’s where only two groups were compared and between groups in One-Way ANOVA’s.

