## Supplementary Figures and Tables for "Selective Vulnerability of Tripartite Synapses in Amyotrophic Lateral Sclerosis"

Latest Edit: 2021/10/28.

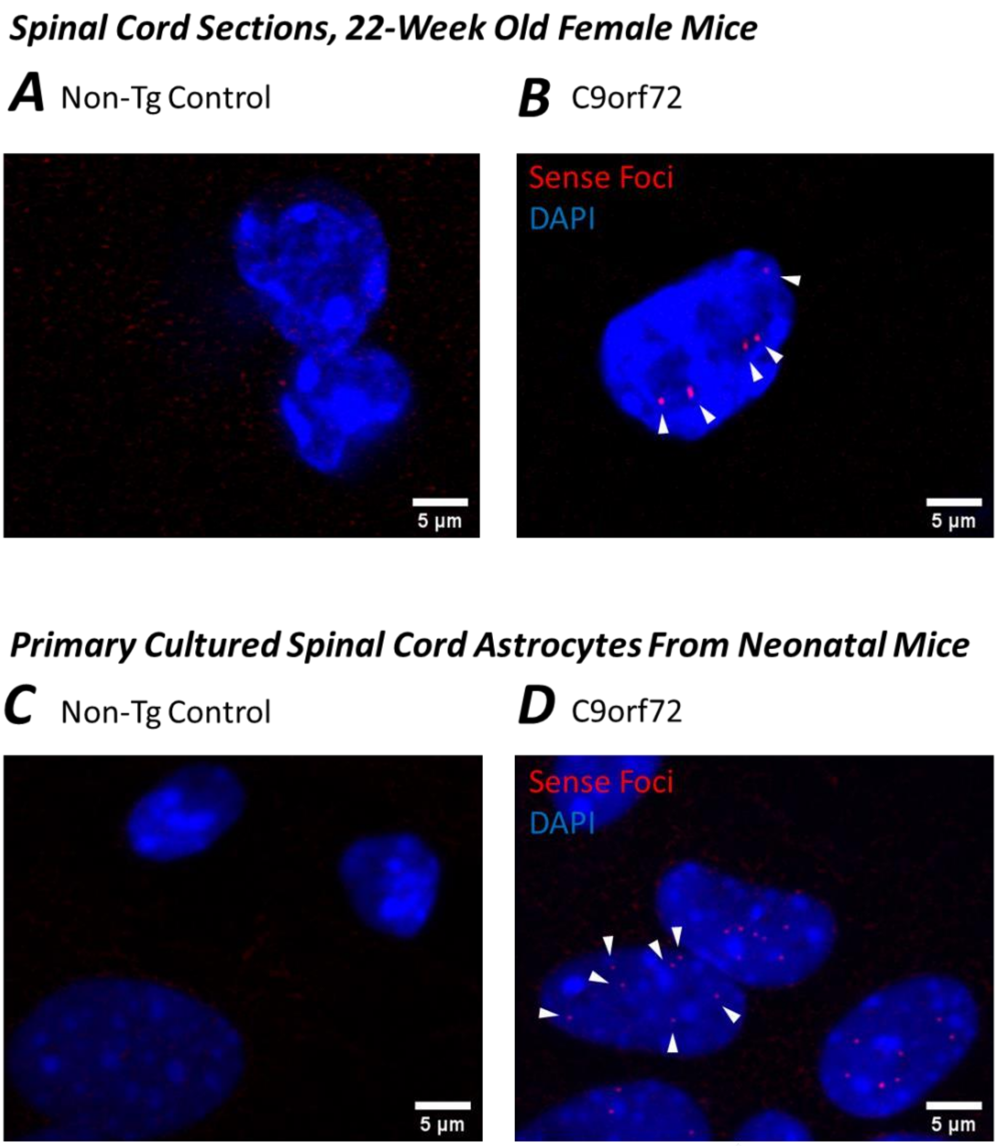


**SI. Figure 1**. **G4C2-repeat RNA foci in C9orf72 mouse spinal cord tissue and primary astrocytes**. Foci are labelled by FISH using Cy5-conjugated RNA probes (red) and appear visible in DAPI-labelled cell nuclei (blue). **A-B**. RNA foci are present in the nuclei of cells in spinal cord sections from a 22-week old female PSD95-eGFP mouse expressing the C9orf72 mutation, but not in an age and gender matched non-Tg control mouse. **C-D**. RNA foci are present in the nuclei of primary cultured astrocytes from C9orf72 neonatal mice, but not in astrocytes cultured from non-Tg littermates.

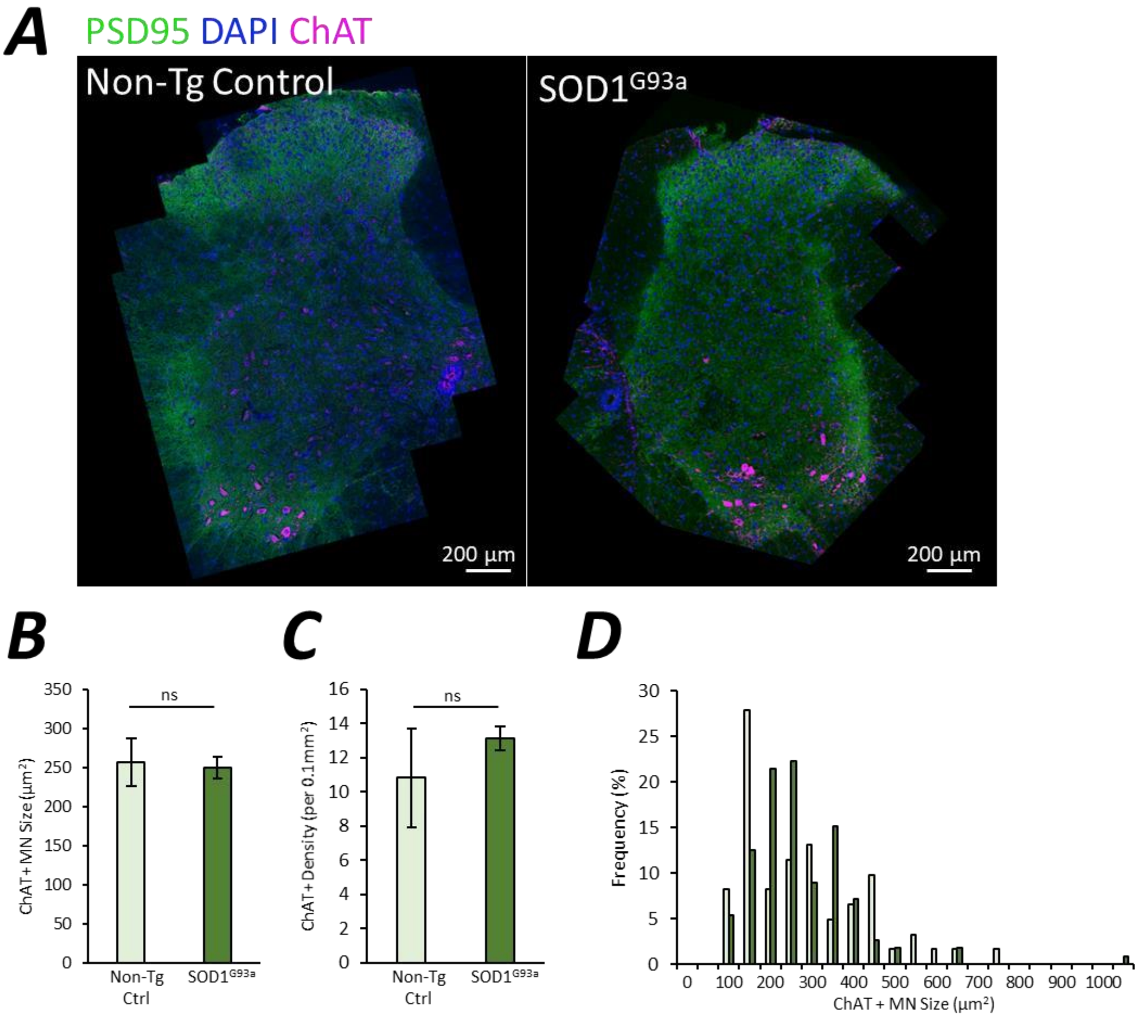


**SI. Figure 2. ChAT immunolabelling of 16 week SOD1$^{G93a}$ mice reveals no significant loss of MNs**. **A**. Example high-resolution maps of PSD95-eGFP, ChAT and DAPI labelling in non-Tg control and SOD1$^{G93a}$ 16 week male spinal cords. **B**. Chart plotting MN size (area) in non-Tg Control and SOD1$^{G93a}$ mice. **C**. Chart plotting MN cell density in non-Tg Control and SOD1$^{G93a}$ mice. **D**. Frequency histogram plotting the number of cells of a given size in non-Tg Control and SOD1$^{G93a}$ mice.

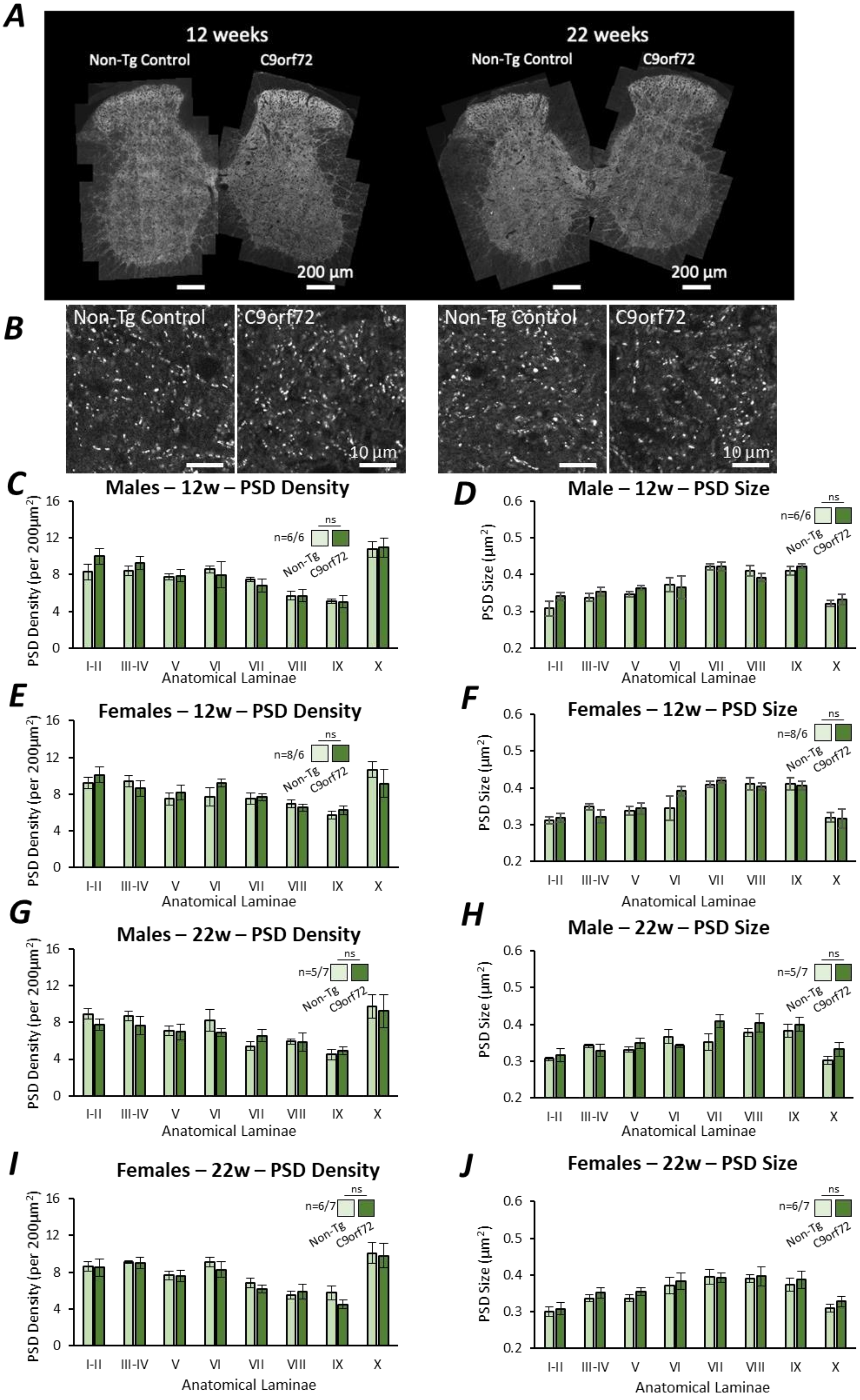
A
12 weeks
Non-Tg Control
C9orf72
22 weeks
Non-Tg Control
C9orf72
200 µm
200 µm
B
Non-Tg Control
C9orf72
10 µm
Non-Tg Control
C9orf72
10 µm
C
Males – 12w – PSD Density
PSD Density (per 200µm²)
n=6/6
ns
Non-Tg
C9orf72
I-II III-IV V VI VII VIII IX X
Anatomical Laminae
D
Male – 12w – PSD Size
PSD Size (µm²)
n=6/6
ns
Anatomical Laminae
E
Females – 12w – PSD Density
n=8/6
F
Females – 12w – PSD Size
n=8/6
G
Males – 22w – PSD Density
n=5/7
H
Male – 22w – PSD Size
n=5/7
I
Females – 22w – PSD Density
n=6/7
J
Females – 22w – PSD Size
n=6/7

**SI. Figure 3. Widescale mapping of excitatory synapses reveals no changes associated with C9orf72 mutation in C9BAC500 mice.** **A.** High-resolution maps of PSD95-eGFP expression in 12 week and 22 week old C9orf72 mice and non-Tg controls. **B**. Example cropped high resolution images of individual PSD's in C9orf72 mice and non-Tg controls. **C**. Chart plotting the PSD density in male 12 week old mice. **D**. Chart plotting the PSD size in male 12 week old mice. **E**. Chart plotting the PSD density in female 12 week old mice. **F**. Chart plotting the PSD size in female 12 week old mice. **G**. Chart plotting the PSD density in male 22 week old mice. **H**. Chart plotting the PSD size in male 22 week old mice. **I.** Chart plotting the PSD density in female 22 week old mice. **J.** Chart plotting the PSD size in female 22 week old mice.

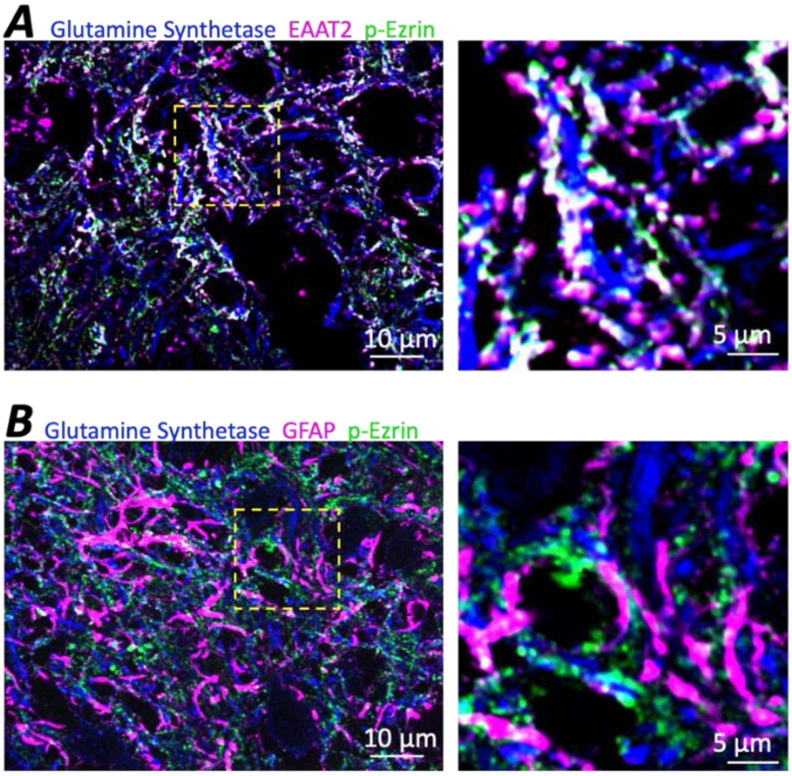


**SI. Figure 4. p-Ezrin and EAAT2 label perisynaptic astrocytic processes in the mouse spinal cord.** Wild-type spinal cord sections from a P90 Wild-type mouse were immunolabelled for astrocytic markers using a combination of conventional primary-secondary antibody labelling and dye-conjugated primary antibody labelling. Glutamine synthetase was visualised with a FITC-conjugated secondary antibody, GFAP and EAAT2 were conjugated directly to Alexa Fluor 594, and p-Ezrin was conjugated directly to Alexa Fluor 647. **A**. p-Ezrin and EAAT closely colocalise along glutamine synthetase-positive astrocyte branches. **B**. p-Ezrin puncta typically associate along glutamine synthetase branches as opposed to GFAP-positive branches, which are more associated with larger astrocytic branches and the soma.

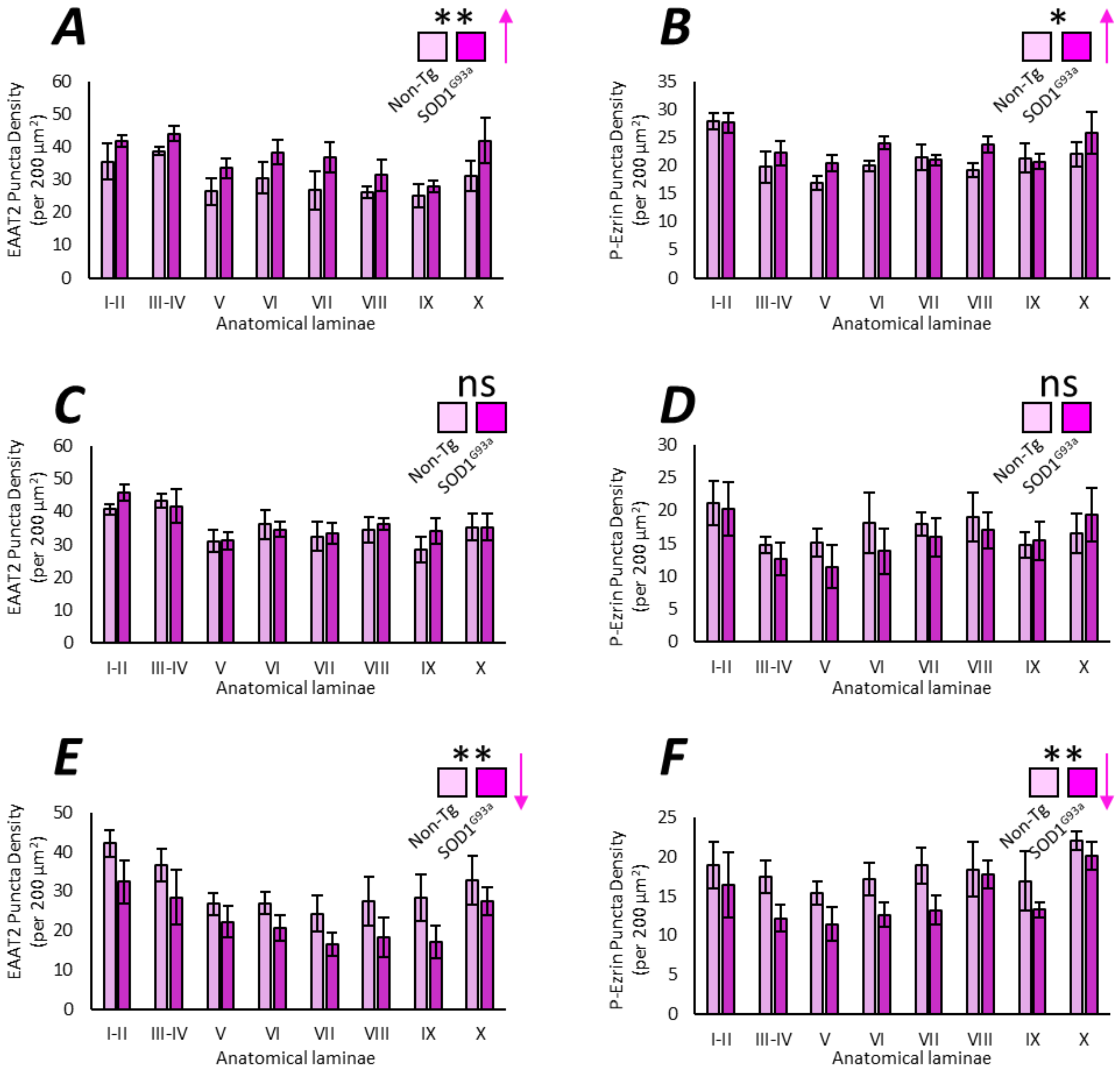


**SI. Figure 5. Mapping PAP marker distribution in the ALS mouse spinal cord reveals age-dependent changes in ALS.** The puncta density of EAAT2 (**A, C, E**) and p-Ezrin (**B, D, F**) were analysed for each spinal cord laminae at 8 weeks (**A-B**), 12 weeks (**C-D**) and 16 weeks (**E-F**). These data reveal increased PAP protein expression at the pre-symptomatic 8 week stage but significantly reduced expression at the early symptomatic 16 week stage.

### Supplementary Tables

| SI. Table 1. Human Post-Mortem Tissue MRC ID's | | | | | |
|---|---|---|---|---|---|
| **MRC Brain Bank ID** | **Sex** | **Age** | **Genotype/Condition** | **Tissue Type** | **Purpose** |
| BBN001.29085 | Male | 46 | Control | Cervical Spinal Cord | Tripartite Synapse Labelling (Fig. 4) |
| BBN001.29084 | Male | 58 | Control | Cervical Spinal Cord | Tripartite Synapse Labelling (Fig. 4) |
| BBN001.26797 | Male | 49 | Control | Cervical Spinal Cord | Tripartite Synapse Labelling (Fig. 4) |
| BBN001.26309 | Male | 69 | Control | Cervical Spinal Cord | Tripartite Synapse Labelling (Fig. 4) |
| BBN001.25751 | Male | 50 | Control | Cervical Spinal Cord | Tripartite Synapse Labelling (Fig. 4) |
| BBN001.29824 | Female | 71 | Control | Cervical Spinal Cord | Tripartite Synapse Labelling (Fig. 4) & IHC Protocol Validation |
| BBN_20613 | Male | 50 | C9orf72 (ALS) | Cervical Spinal Cord | Tripartite Synapse Labelling (Fig. 4) |
| BBN_20993 | Male | 43 | C9orf72 (ALS) | Cervical Spinal Cord | Tripartite Synapse Labelling (Fig. 4) |
| BBN001.28792 | Male | 58 | C9orf72 (ALS) | Cervical Spinal Cord | Tripartite Synapse Labelling (Fig. 4) |
| BBN001.35827 | Male | 66 | C9orf72 (ALS) | Cervical Spinal Cord | Tripartite Synapse Labelling (Fig. 4) |
| BBN001.35506 | Male | 71 | C9orf72 (ALS) | Cervical Spinal Cord | Tripartite Synapse Labelling (Fig. 4) |
| BBN001.36134 | Male | 45 | SOD1 (ALS) | Cervical Spinal Cord | Tripartite Synapse Labelling (Fig. 4) |
| BBN001.35135 | Male | 71 | SOD1 (ALS) | Cervical Spinal Cord | Tripartite Synapse Labelling (Fig. 4) |
| BBN001.34243 | Male | 53 | SOD1 (ALS) | Cervical Spinal Cord | Tripartite Synapse Labelling (Fig. 4) |
| BBN001.29543 | Male | 47 | SOD1 (ALS) | Cervical Spinal Cord | Tripartite Synapse Labelling (Fig. 4) |
| BBN00135209 | Female | 70 | SOD1 (ALS) | Lumbar Spinal Cord | IHC Protocol Validation |
| BBN001.29825 | Female | 70 | C9orf72 (ALS) | Lumbar Spinal Cord | IHC Protocol Validation |
